# From prototype to outbreak: conserved pathogenesis of Oropouche virus in a novel murine pregnancy model highlights its public health implications

**DOI:** 10.1101/2025.08.02.668287

**Authors:** Krista B. Gunter, James M. Bowen, Andrew T. Clarke, Melanie McFarlane, Dorcus C. A. Omoga, Stephanie Pozuelos, Lisa M. Rogers, David M. Aronoff, Jay Vornhagen, Benjamin Brennan, Natasha L. Tilston

## Abstract

Oropouche virus (OROV) is an emerging orthobunyavirus responsible for widespread outbreaks across South and Central America. The recent surge in congenital infections has raised urgent concerns about OROV’s threat to maternal and fetal health. Here, we establish an *in vivo* model of OROV vertical transmission using the ancestral (prototype) strain BeAn19991 in immunocompetent C57BL/6J mice. We demonstrate that OROV efficiently replicates in maternal tissues, crosses the maternal–fetal interface, and infects both placental and fetal tissues. Parallel infections in human trophoblast-derived cell lines confirm conserved placental tropism across the ancestral strain and a contemporary (outbreak) isolate from the current outbreak. Importantly, we show that vertical transmission is not a recently acquired trait but a long-standing feature of OROV biology. Offspring born to infected dams mount neutralizing antibody responses and exhibit partial protection upon challenge. These findings conclusively confirm OROV as a vertically transmissible arbovirus, highlighting the urgent need to integrate OROV into surveillance, diagnostic, and vaccine preparedness efforts.

## Introduction

As of mid-2025, Oropouche fever cases continue to rise across the Americas, with the highest case counts reported in Brazil, Cuba, and Panama. Since the onset of the outbreak in late 2022, over 23,000 confirmed cases have been documented, with Brazil accounting for the majority^1–3^. A marked surge in late 2023 prompted the Pan American Health Organization/World Health Organization (PAHO/WHO) to issue an epidemiological alert, underscoring the virus’s expanding public health threat^4–10^. Oropouche virus (OROV), the etiologic agent of Oropouche fever, is an orthobunyavirus primarily transmitted by *Culicoides paraensis* midges and has circulated endemically in parts of South America since the 1960s^11^. However, the current outbreak is unprecedented in both scale and geographic distribution. For the first time, OROV has spread beyond its traditional range into new countries such as Cuba, which reported 626 confirmed and over 24,000 suspected cases in 2024 alone^12^. This expansion has resulted in imported cases in non-endemic regions, including the United States, Canada, the United Kingdom, and continental Europe^1^. Notably, we identified *Culicoides sonorensis*, a widespread North American midge species, as a competent vector for OROV^13^, raising the possibility of local transmission if the virus is introduced. Combined with the accelerating effects of climate change on vector ecology^14,15^, these findings raise concerns about future OROV establishment in North America.

In parallel with its geographic spread, OROV’s clinical profile appears to be evolving, likely driven not only by viral genetic changes but also by rising case numbers associated with shifting human behaviors, such as increased urbanization. Once considered a self-limiting febrile illness, OROV is now associated with severe neurological complications^1^, mother-to-fetus transmission^16–18^, and fatalities^1,19^, clinical outcomes not previously reported at this scale. These developments highlight the growing public health threat posed by historically overlooked arboviruses and underscore the urgent need to reassess OROV’s pathogenic potential and epidemic capacity.

OROV is a tri-segmented, single-stranded, negative-sense RNA virus (*Orthobunyavirus oropoucheense*, *Peribunyaviridae*, *Bunyavirales*)^20^. Its genome comprises three segments: the S segment encodes the nucleoprotein (N) and the non-structural protein NSs in overlapping open reading frames; the M segment encodes a glycoprotein precursor that is co-translationally cleaved into glycoproteins Gn and Gc, as well as the NSm protein; and the L segment encodes the RNA-dependent RNA polymerase (RdRp). Each segment is flanked by untranslated regions (UTRs) that harbor signals essential for replication, transcription, and packaging^21^. OROV’s segmented genome facilitates reassortment, a process in which genome segments are exchanged during co-infection with other orthobunyaviruses, thereby accelerating evolutionary change^11,22^. Multiple naturally occurring reassortants, such as Iquitos virus, Madre de Dios virus, and Perdões virus, have emerged through inter-species reassortment events, often involving an unidentified donor of the M segment^23^. Although rare, such events can result in substantial shifts in virulence or host range^24,25^. In contrast, intra-species reassortment among circulating OROV strains appears to be more frequent and has been implicated in the diversification of strains associated with the current outbreak^3^.

While intra-species reassortment is often phenotypically neutral, the unprecedented rise in congenital infections during the recent outbreak has raised concern that reassortment contributed to changes in viral tropism, immune evasion, and pathogenesis^1^. The extent to which genome segment exchange influences OROV virulence, particularly in the context of maternal-fetal transmission, remains poorly understood and represents a critical gap in our knowledge. Here, we developed a novel murine model of OROV infection during pregnancy using the ancestral strain BeAn19991 to investigate the virus’s capacity for vertical transmission. Through a combination of murine gestational infections and parallel studies in human placental cell lines, we demonstrate that OROV readily crosses the maternal–fetal interface and infects placental and fetal tissues. Importantly, our findings reveal that vertical transmission is not a novel feature acquired by contemporary isolates but rather a conserved property of OROV biology that has likely been historically overlooked. These results not only establish the first immunocompetent model of congenital OROV infection but also provide critical experimental evidence that reframes our understanding of OROV pathogenesis and underscores its potential as a congenital arboviral threat.

## Results

### OROV replicates efficiently in the liver and spleen of immunocompetent C57BL/6J mice

To establish a physiologically relevant model of OROV infection during pregnancy, we first assessed whether immunocompetent wild-type (WT) C57BL/6J mice support viral replication. Using our previously validated recombinant (r) reporter OROV (rOROVMZsG)^26^, which we demonstrated was lethal in type I interferon (IFN) receptor knock-out (IFNAR^-/-^) mice, we conducted a 14-day infection study (Supplemental Figure 1a). Infected mice exhibited no overt clinical signs and gained weight comparably to animals inoculated with UV-inactivated virus, indicating minimal disease burden (Supplemental Figure 1b). Quantification of viral RNA (vRNA) across tissues revealed OROV replication in the liver and spleen at 5- and 7-days post-infection (dpi). In contrast, vRNA levels in the heart, lung, and brain were considerably lower or not detectable (ND) (Supplemental Figure 1c). By 14 dpi, vRNA loads had declined, consistent with viral clearance. Infectious virus was isolated from liver and spleen homogenates at 5 dpi, confirming active replication (Supplemental Figure 1d). Collectively, these results demonstrate that rOROV establishes a productive, self-limiting infection in WT C57BL/6J mice, with primary replication localized to the liver and spleen.

### OROV infects the placenta and crosses the maternal–fetal barrier in C57BL/6J mice

Next, to investigate the potential for vertical transmission, we subcutaneously inoculated timed-pregnant C57BL/6J dams with rOROV ancestral strain BeAn19991^27^ at embryonic day (E) 4.5-7.5 or E12.5, corresponding to early and mid-gestation, respectively (Fig. 1a). In both groups, dams gained weight steadily throughout pregnancy, with no evidence of fetal reabsorption or pregnancy loss (Fig. 1b, e). Maternal vRNA loads in both groups revealed high viral burdens in the liver and spleen, with mid-gestation infections showing higher levels than early gestation (Fig. 1c, f). This is consistent with (Supplemental Figure 1c), where peak OROV replication occurs at ∼5 dpi, and begins to decline by day 7, corresponding to the mid-gestation cohort being harvested at 5 dpi, and the early cohort at 8 dpi. To evaluate vertical transmission, we quantified vRNA in matched placentas (circles) and fetuses (squares) collected at necropsy. OROV vRNA in placental tissue was consistently detected in both early and mid-gestation groups, though levels were highest following mid-gestation infection (Fig. 1g). Fetal tissues from both groups also harbored detectable vRNA, but at lower levels and with greater variability (Fig. 1d, g). Gross fetal morphology appeared normal across all groups, with no visible developmental abnormalities observed at either gestational stage (Supplemental Figure 2a).

**Fig. 1.**
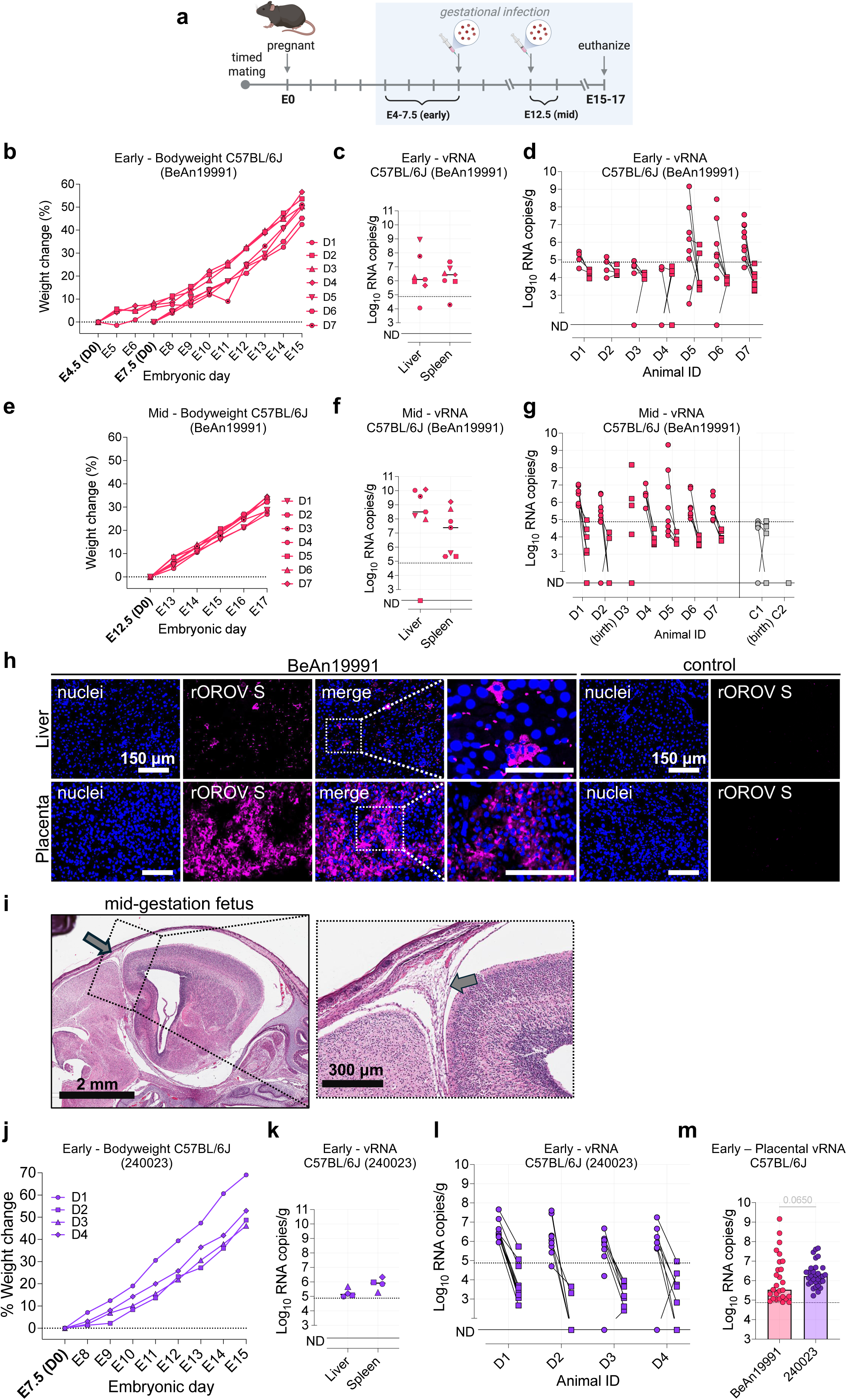
Infection with OROV leads to systemic maternal infection and placental dissemination. (a) Schematic of experimental design. Six-week-old C57BL/6J female mice were housed with C57BL/6J male mice for 48 h to permit mating. Pregnant dams were infected SC with BeAn19991 or Opti-MEM at E4-7.5 (early gestation), or E12.5 (mid-gestation), and euthanized between E15-17. (b, e) Percent weight gain of pregnant BeAn19991-infected dams following early gestation (b) or mid (e) gestation. (c, f) vRNA levels in maternal liver and spleen at harvest, measured by RT-qPCR at early (c) or mid (f) gestation. (d, g) vRNA levels in matched placentas (circles) and fetuses (squares) from each dam, quantified by RT-qPCR at early (d) or mid (g) gestation. (h) HCR RNA-FISH detection of rOROV S segment vRNA (magenta) in maternal liver and placental sections from BeAn19991-infected dams. Infected hepatocytes (liver; top) exhibit cytoplasmic or perinuclear vRNA accumulation. Placental sections (bottom) reveal widespread infection in the labyrinth zone. Nuclei stained with Hoechst (blue). (i) Representative BeAn19991 infected mid-gestation fetus showing focal lymphocytic infiltration in the brain. Enlarged view shown at the side. (j) Percent weight gain of pregnant 240023-infected dams following early gestation infection. (k) vRNA levels in the liver and spleen of pregnant 240023-infected dams at early gestation (l) vRNA levels in matched placentas (circles) and fetuses (squares) from each 240023-infected dam, quantified by RT-qPCR at early gestation. (m) Early gestation placental vRNA loads in mice infected with rOROV BeAn19991 or OROV 240023. Each point represents an individual placenta. The dashed line represent the limit of detection, ND = not detected.

To visualize vRNA, we applied a custom hybridization chain reaction (HCR) RNA-FISH assay targeting OROV S vRNA, as described in our recent study^28^. In infected liver sections, we detected discrete foci of vRNA, with signal localized to the cytoplasm or perinuclear regions of hepatocytes, consistent with active viral replication (Fig. 1h). No vRNA signal was detected in liver sections from control animals. In placental sections from infected dams, widespread vRNA accumulation was observed throughout the labyrinth zone, a key site of maternal–fetal exchange, indicating robust replication at the maternal–fetal interface (Fig. 1h). In contrast, no signal was detected in placentas from mock-infected controls. The distribution of vRNA signal in maternal liver and the placental tissues indicate localized clusters of infection, consistent with productive viral replication. To confirm that vRNA detection reflected infectious virus, placental homogenates were used to inoculate Vero E6 cells. Immunofluorescence imaging at 24 hours post-infection (hpi) revealed rOROV-positive foci in multiple samples (Supplemental Figure 2b), confirming the presence of replication-competent virus. Histopathological analysis of maternal livers revealed mild to moderate extramedullary hematopoiesis (EMH) in rOROV-infected dams compared to mock-infected controls, consistent with localized inflammation or early-stage pathology (Supplemental Figure 2c). Examination of fetal tissues infected at mid-gestation revealed vacuolation of the brains in several fetuses, with one displaying focal lymphocytic infiltrate (Fig. 1i). These findings are indicative of viral invasion, consistent with patterns observed in other congenital viral infections^29^. Taken together, these data demonstrate that the ancestral OROV strain BeAn19991 productively replicates in the placenta following both early and mid-gestational infection. While fetal infection was less frequent, the detection of vRNA in select fetal tissues and replication-competent virus in placentas confirms that BeAn19991 can breach the maternal–fetal barrier and establish an infection at the maternal–fetal interface.

Next, we evaluated whether a contemporary OROV isolate from the current outbreak crosses the maternal–fetal barrier more effectively than the ancestral strain. OROV strain 240023, obtained from the World Reference Center for Emerging Viruses and Arboviruses (WRCEVA) and originally isolated in July 2024 from a returning traveler from Cuba, was used to assess vertical transmission following early gestation infection. All dams gained weight steadily throughout pregnancy with no evidence of fetal loss (Fig. 1j). vRNA was readily detected in maternal liver and spleen (Fig. 1k), and placental viral loads (Fig. 1l, circles) consistently exceeded those in matched fetal tissues (Fig. 1l, squares), mirroring patterns observed with rOROV BeAn19991. The frequency of placental infection was significantly higher at early gestation in dams infected with OROV 240023 compared to the BeAn19991 group (88.57% vs. 57.45%, χ² test, *p* = 0.0022). Moreover, early gestation infection placental vRNA levels in the 240023-group trended higher than those in the BeAn19991 group (Mann–Whitney test, *p* = 0.0650) (Fig. 1m). We also assessed mid gestation (E14.5) with OROV 240023. High vRNA was detected in maternal (Supplemental Figure 2d) and corresponding placental (Supplemental Figure 2e) tissues of animals D1 and D5, while animals D2 and D6 in this group had progressed to delivery. Collectively, these results demonstrate that OROV 240023 efficiently infects the placenta and crosses the maternal–fetal barrier in immunocompetent mice, with replication dynamics trending potentially better than the ancestral strain.

To explore whether this observation reflected an enhanced replicative fitness in human placental tissues, we used three trophoblast-derived cell lines. These *in vitro* systems allowed us to directly assess the conservation of placental tropism across species and determine the human relevance of our murine findings. We compared rOROV BeAn19991 and OROV 240023 in HTR8 (SV40-transformed first-trimester extravillous trophoblasts), BeWo (choriocarcinoma), and JEG3 (choriocarcinoma) cells. Both strains productively infected all three lines, with replication consistently highest in HTR8 cells, mirroring the preferential infection observed in first-trimester placentas in our mouse model. By 72 hpi, both viruses achieved comparable peak titers (Fig. 2a–c). Immunofluorescence imaging confirmed widespread infection and cytoplasmic accumulation of viral antigen in each model, with no discernible differences in spread or cytopathic effect (Fig. 2d–g). Together, these findings confirm that OROV is intrinsically capable of infecting placental cells and crossing the maternal–fetal barrier, a conserved feature of its biology independent of strain.

**Fig. 2.**
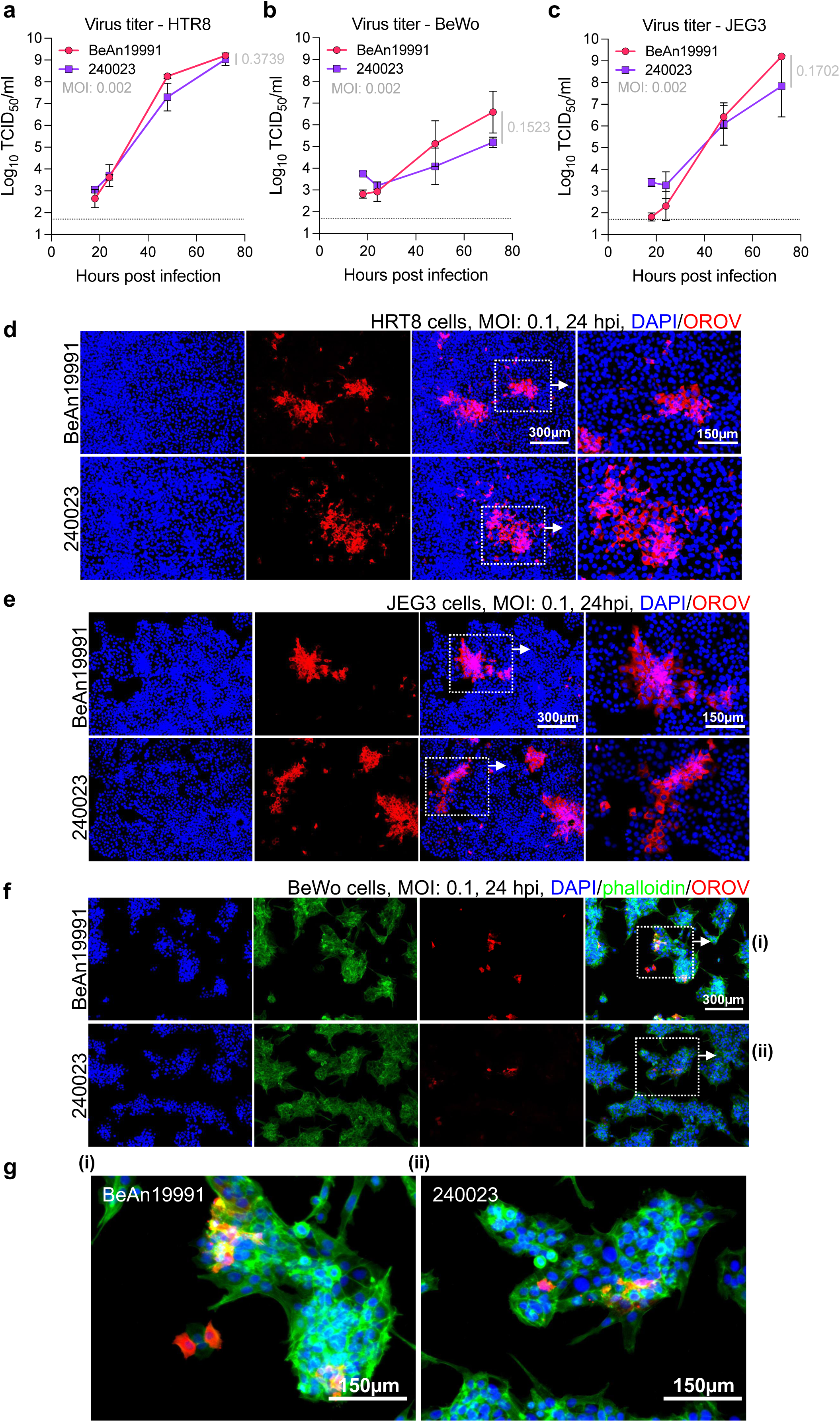
Ancestral and contemporary OROV replicate efficiently in human placental-derived cell lines. (a–c) Growth kinetics of rOROV BeAn19991 and OROV 240023 in human placental cell lines: HTR8 (a), BeWo (b), and JEG3 (c). Cells were infected at MOI 0.002, and supernatants were collected at 18, 24, 48, and 72 hpi. Each point represents an independent replicate; bars indicate mean ± SD. The p-value was calculated for the 72-h time point. (d–g) Immunofluorescence imaging of infections in cells at MOI 0.1, stained with anti-OROV antibody (red) and DAPI (blue). (g) High magnification fields of (f) Scale bars as indicated. BeWo cells were also stained with phalloidin (green). (EVOS M5000 imaging system, ThermoFisher).

### Contemporary OROV isolates exhibit enhanced replication relative to the ancestral strain

To compare phenotypic differences between the ancestral and contemporary OROV, we compared their replication in both type I IFN-competent (A549) and IFN-deficient (Vero E6) cells. Although both viruses reached similar titers by 48 hpi, OROV 240023 consistently exhibited approximately one log higher titers at 24 hpi in both cell types (Fig. 3a, *p* = 0.0021 in A549 cells, *p* = 0.0020 in Vero E6), indicating a replicative advantage during early infection.

**Fig. 3.**
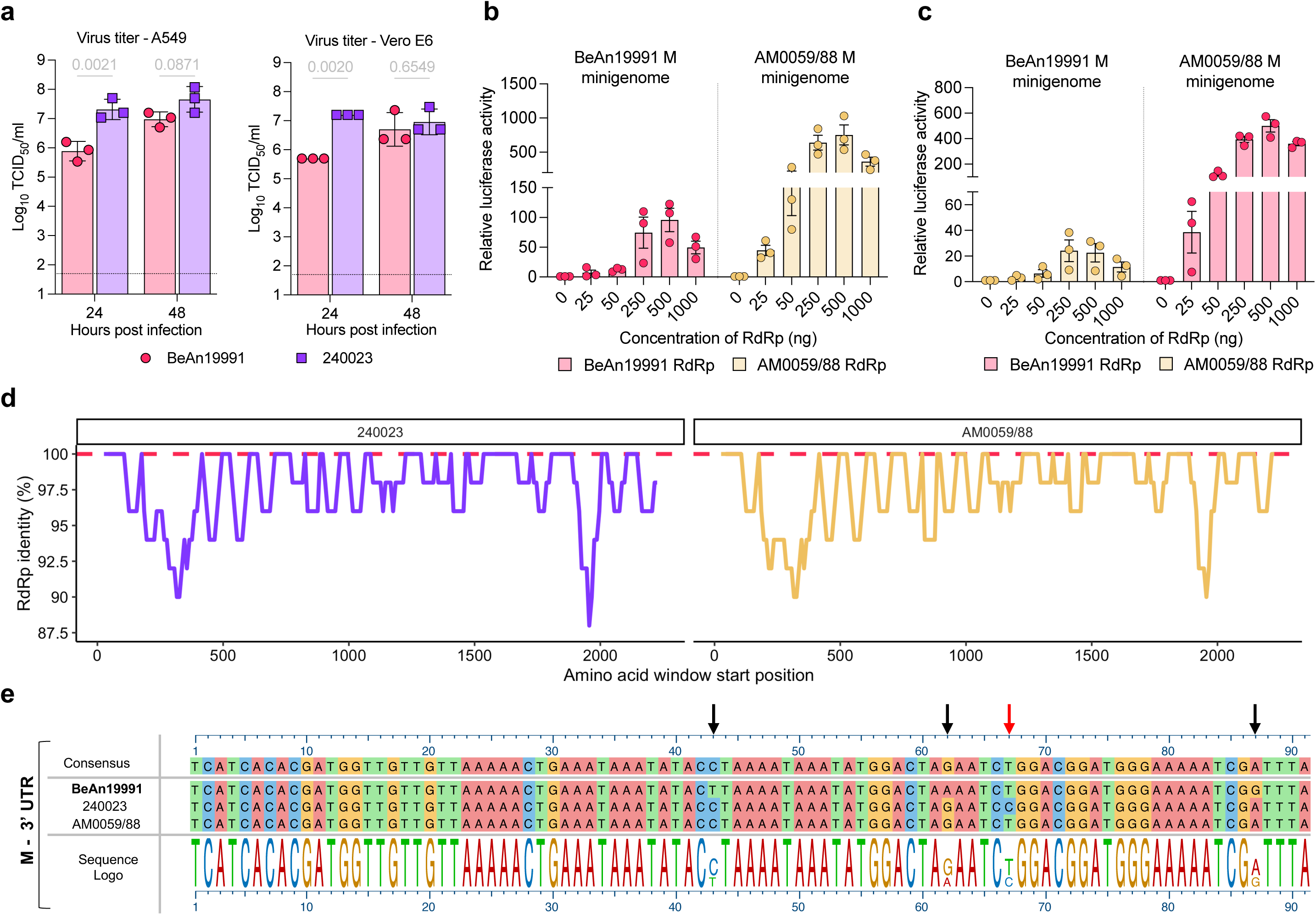
Contemporary OROV strains exhibit enhanced growth and altered polymerase activity. (a) Growth kinetics of rOROV BeAn19991 and OROV 240023. Supernatants were collected at 24 and 48 hpi (MOI 0.1) and titrated by TCID_50_ assay. Each point represents an individual replicate; bars show mean ± SD. (b, c) Cross-comparison of polymerase activity using BeAn19991 or AM0059/88 RdRp in M segment minigenome systems from BeAn19991 (red) and AM0059/88 (yellow). BSRT7/5 cells were transfected with an optimized ratio of N and RdRp along with the M segment minigenome (Supplemental Figure 3, and^35^). Luciferase activity was measured 24 hpt. (d) Sliding window analysis of amino acid identity across the RdRp of OROV 240023 (purple) and AM0059/88 (yellow) relative to BeAn19991 (dashed red). Identity values were plotted along the RdRp using a 50-amino-acid sliding window. (e) Alignment of the M segment 3′ UTR from BeAn19991, 240023, and AM0059/88. Black arrows indicate substitutions between 240023, AM0059/88 vs BeAn19991, and the red arrow denotes a unique insertion in 240023. The sequence logo below shows nucleotide conservation.

As an additional point of comparison, we incorporated data for the Brazilian isolates OROV AM0088 and AM0059 (treated together as AM0059/88; isolated in Manaus, Brazil, January 2024). In our study, we treat OROV AM0059/88 and 240023 as representative contemporary isolates, enabling direct comparison with BeAn19991. Interestingly, Scachetti *et al.*^30^ had previously reported a replication advantage for OROV AM0088 compared to BeAn19991. However, in their system, BeAn19991 failed to reach comparable titers in any of the tested cell lines. As we were unable to obtain live Brazilian isolates for side-by-side testing, we instead received their full-genome sequences (Drs. William M. de Souza, José Luiz Proenca-Modena, and Pritesh Lalwani; GenBank accession numbers PP992525–PP992530). To assess transcriptional/replication efficiency under our experimental conditions, we employed a minigenome system as a surrogate. Among the three segments, only the M segment contained complete 3′ and 5′ UTRs, and we therefore generated an M–segment–based minigenome (AM0059/88 M) encoding a humanized *Renilla* luciferase reporter. Luciferase activity from the AM0059/88 minigenome was robust and increased in a dose-dependent manner with both N and RdRp expression, peaking at a 1:1 ratio (Supplemental Figure 3). Compared to the BeAn19991 M minigenome, AM0059/88 consistently exhibited ∼10-fold higher luciferase activity (Fig. 3b, c). These results indicate that the AM0059/88 M segment confers enhanced transcriptional/replication efficiency even in the absence of a cognate RdRp, suggesting that its UTR regulatory elements may drive increased activity independently.

To identify potential molecular determinants underlying this effect, we performed sliding-window analysis of the RdRp amino acid sequence. The polymerase of OROV 240023 and AM0059/88 displayed multiple regions of divergence relative to BeAn19991, particularly in the N- and C-terminal regions (Fig. 3d). While the N protein was conserved entirely across all three strains, these RdRp differences may contribute to enhanced polymerase processivity or stability. In addition, we noted three shared nucleotide differences in the M segment 3′ UTR of OROV 240023 and AM0059/88 that were absent in BeAn19991 (Fig. 5e). The 5′ UTR was fully conserved across all strains, suggesting that these 3′ UTR polymorphisms may influence replication or transcriptional efficiency through altered secondary structure or interaction with the polymerase complex. Together, these data reveal functional differences between ancestral and contemporary OROV isolates at both the viral replication and polymerase levels, with implications for viral fitness.

### Contemporary OROV isolate exhibits enhanced virulence, immune escape, and sequence divergence from the ancestral strain

To evaluate the *in vivo* pathogenicity of contemporary OROV isolate 240023, we used our validated lethal IFNAR^⁻/⁻^ mouse model^26^. Mice infected with OROV 240023 succumbed to disease earlier than those infected with the ancestral strain, with most succumbing to primary infection. However, one survivor was observed in the OROV 240023 group and two in the BeAn19991 group, derived from two independent experiments (Fig. 4a). For the BeAn19991 group, only one of the two survivors was re-challenged. Upon homologous re-challenge at 14 dpi, both survivors succumbed to disease by 4 dpi. Clinical symptoms, including ruffled fur, hunched posture, and weakness, were noted in both groups (Fig. 4b), and pale livers were observed at necropsy in several animals. Both groups also exhibited rapid weight loss consistent with what we have previously noted with rOROV BeAn19991 in IFNAR^⁻/⁻^ mice^26^ (Fig. 4c, d). High vRNA loads were detected in all major organs, including liver, spleen, heart, lung, and brain, confirming widespread dissemination (Fig. 4e, f) and likely contributing to the more rapid disease progression observed in OROV 240023-infected mice.

**Fig. 4.**
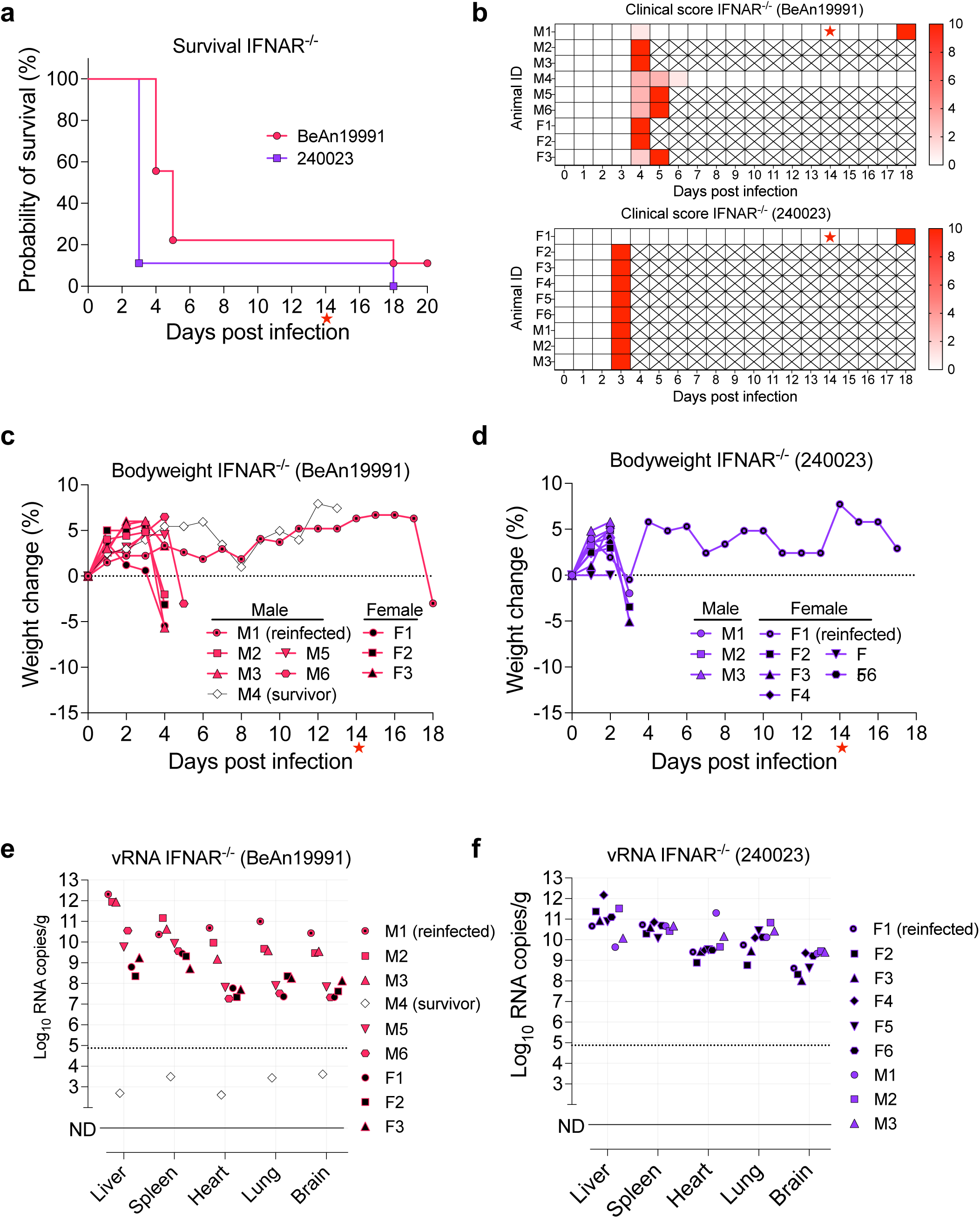
Contemporary OROV strain 240023 induces earlier mortality and higher viral burden in IFNAR^⁻/⁻^ mice. (a) Kaplan–Meier survival curves for mice infected with rOROV BeAn19991 or OROV 240023. (b) Clinical disease scoring matrix for each animal, with dpi shown on the x-axis and cumulative clinical scores indicated by heatmap scale (red = higher severity; X = euthanasia). (c, d) Percent weight change over time in individual male (M) and female (F) mice infected with BeAn19991 (c) or 240023 (d). (e, f) vRNA loads in liver, spleen, heart, lung, and brain tissues of infected mice, measured by RT-qPCR at euthanasia. Data are shown for individual mice infected with rOROV BeAn19991 (e) or OROV 240023 (f). Mice that were reinfected or survived to the study endpoint are indicated. Red stars indicate the time of rechallenge for survivors.

Given its enhanced virulence, we next examined the antigenic similarity of OROV 240023 to the ancestral strain. Virus neutralization tests (VNT) using serum from rOROV BeAn19991-infected mice revealed a reduction in cross-neutralization of OROV 240023, despite robust neutralization of the homologous strain (Fig. 5a, b), suggesting antigenic divergence and potential immune escape. To investigate the molecular basis of this finding, we compared amino acid sequences across the M segment, which encodes the surface glycoproteins (Gn/Gc) and the NSm protein. OROV 240023 and AM0059/88 shared multiple nonsynonymous substitutions relative to the ancestral strain (Fig. 5c). In agreement with prior studies on the Brazilian isolate AM0088, differences in the Gc likely contribute to the observed reduction in cross-neutralization^30^. In contrast to AM0088^30^, OROV 240023 had a non-significant difference in plaque morphology compared to rOROV BeAn19991 [unpaired t-test, *p*= 0.0818; mean plaque diameters: 2.4 mm (BeAn19991) vs 2.7 mm for (240023)] (Fig. 5d). In parallel, sequence alignment of the NSs protein, a known IFN antagonist, revealed differences between the three strains, including conserved substitutions in both OROV 240023 and AM0059/88 and one unique variation in OROV 240023 only (Fig. 5e, red arrow). Together, these data demonstrate that contemporary OROV isolates not only replicate more efficiently *in vitro* but also exhibit enhanced virulence *in vivo*, reduced susceptibility to ancestral-specific neutralizing antibodies, and divergence across multiple proteins, reinforcing the need for in-depth virus-host interaction studies.

**Fig. 5.**
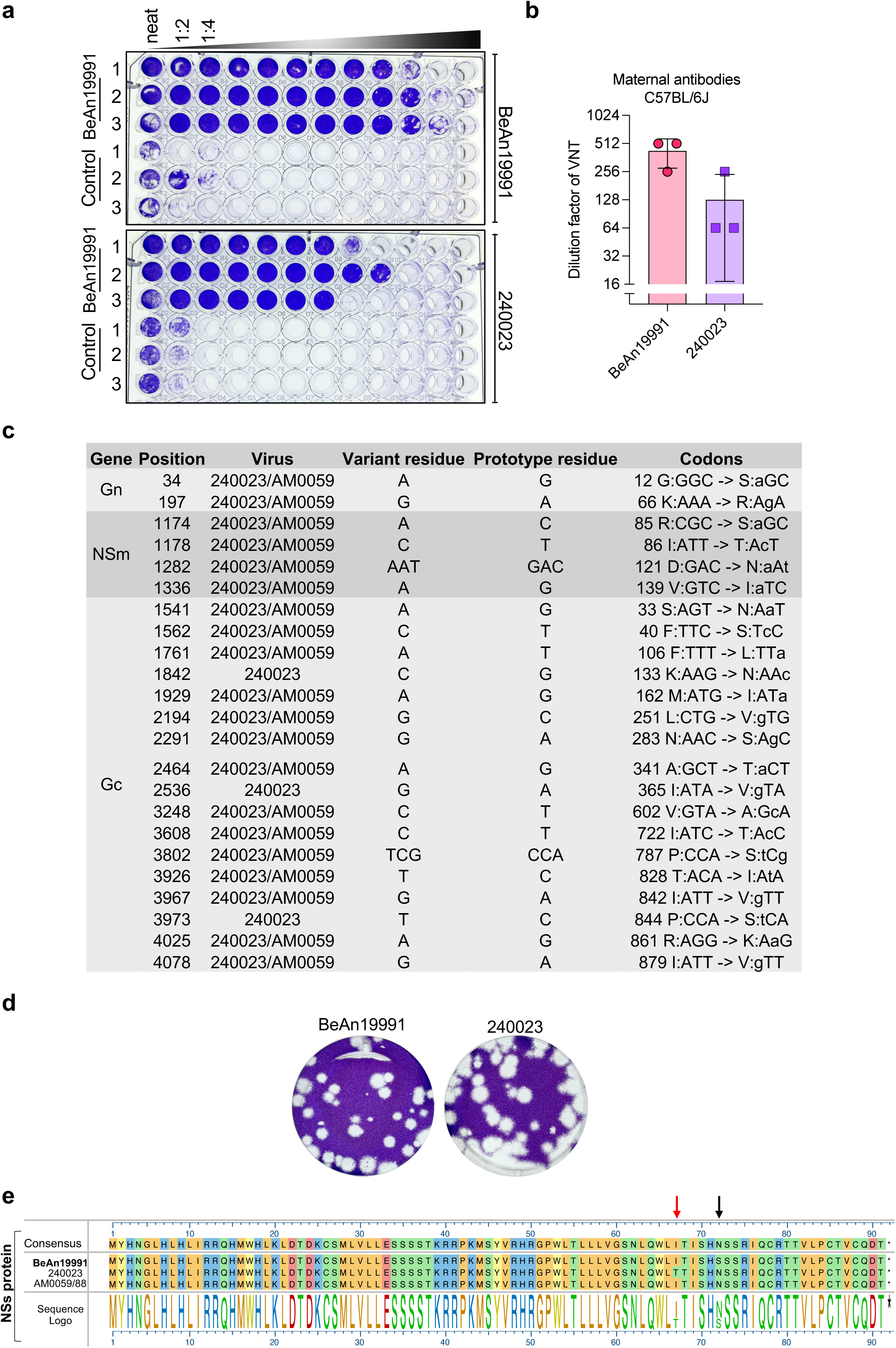
Contemporary OROV isolate 240023 shows reduced neutralization sensitivity and sequence divergence in structural and non-structural proteins. (a) Representative VNT plates showing serial 1:2 serum dilutions (first column = highest concentration) from mice infected with rOROV BeAn19991. Vero E6 cells were fixed and stained with crystal violet at 5 dpi (b) VNT titers for rOROV BeAn19991 and OROV 240023. Each point represents an individual animal; bars indicate mean ± SD. (c) Table of nonsynonymous coding changes in the M segment (Gn, NSm, Gc) of OROV 240023 and/or AM0059/88 compared to BeAn19991. Codon changes and residue substitutions are shown. (d) Plaque morphology of rOROV BeAn19991 and OROV 240023 on Vero E6 cells, stained with crystal violet. (e) Amino acid alignment of the NSs protein from BeAn19991, 240023, and AM0059/88. A sequence logo below highlights conserved and variable residues. Black and red arrows denote polymorphisms of interest, including a unique substitution in 240023 (red arrow).

### Maternal infection with rOROV BeAn19991 confers partial protection to offspring through the production of neutralizing antibodies

Given OROV’s ability to cross the maternal–fetal barrier and its potential for immune evasion, we next investigated whether maternal infection could influence the susceptibility of offspring to subsequent infection. Specifically, we tested whether infection during pregnancy with the prototype strain rOROV BeAn19991 conferred protective immunity to pups following post-weaning challenge. To do this, we challenged pups born to mock-infected (naive) or rOROV-infected (exposed) C57BL/6J dams at 3 weeks of age and monitored infection outcome over two weeks (Fig. 6a). vRNA levels in the liver, spleen, and brain were measured at 3, 5, 7, 10, and 14 dpi. Pups born to naive dams exhibited high levels of vRNA across all tissues, with peak replication at 7–10 dpi (Fig. 6b–d). Notably, one pup succumbed to disease at 7 dpi with high levels of vRNA in the brain. Pups also exhibited signs of disease at 10 dpi with hunched posture and ruffled fur. In contrast, pups born to rOROV-exposed dams exhibited markedly reduced vRNA levels at all time points and no clinical symptoms, consistent with passive protection mediated by maternally transferred immunity.

**Fig. 6.**
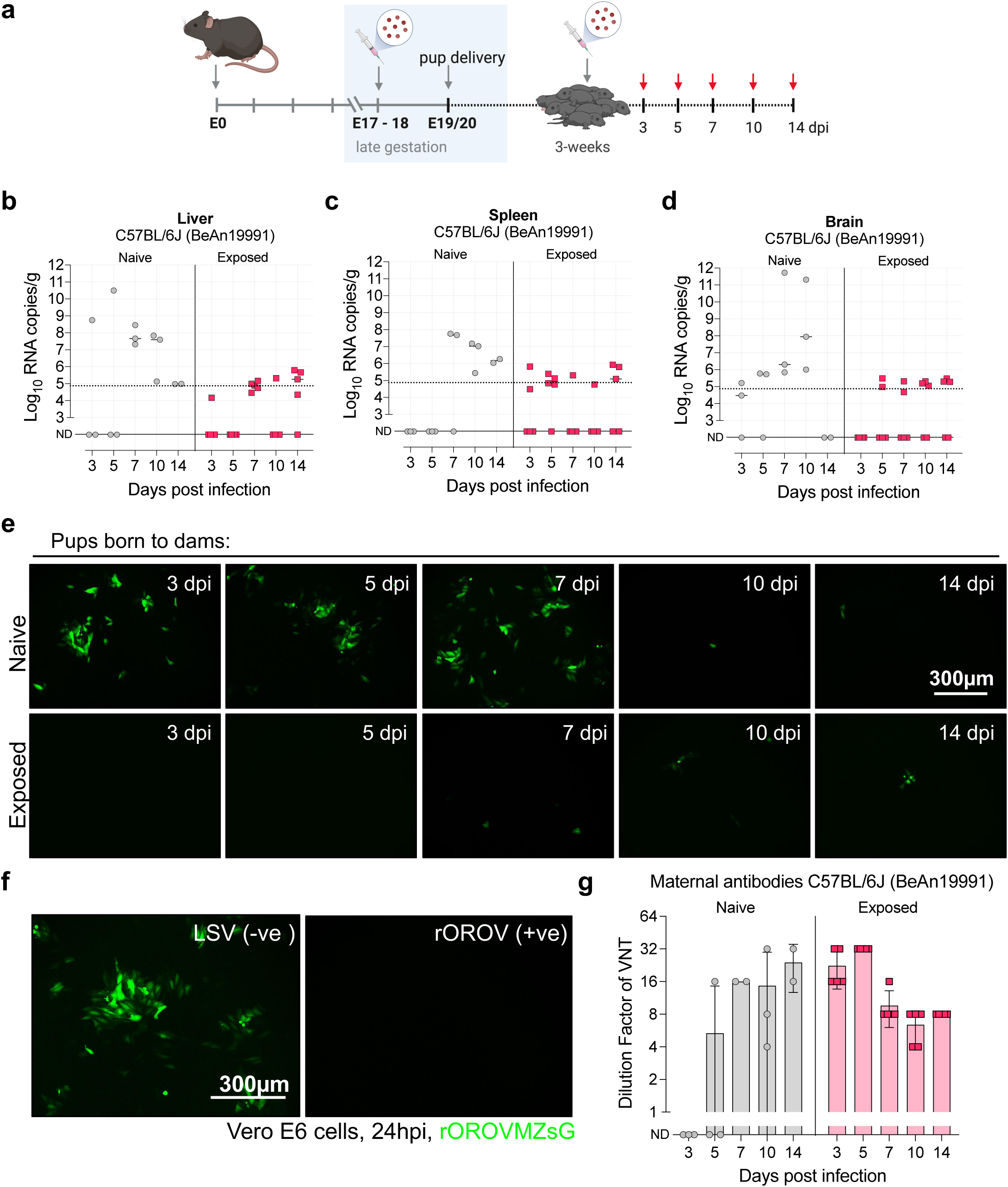
Maternal rOROV BeAn19991 infection confers partial protection to offspring via neutralizing antibodies. (a) Schematic of experimental design. (b-d) Viral load per gram of tissue in the liver (b), spleen (c), and brain (d) of pups, as measured by RT-qPCR. Pups born to rOROV-exposed dams (red squares). Pups born to rOROV naive dams (grey circles). (e) Representative images of rOROVMZsG-based VNT in Vero E6 cells using serum collected from challenged pups. Fluorescence microscopy at 24 hpi shows ZsGreen signal (green) in the presence of serum from pups born to naive or exposed dams across timepoints. (f) Quantification of neutralization titers from fluorescence readout. Dilution factors indicate peak neutralizing activity at early timepoints, with higher titers in pups born to exposed dams.

To assess neutralizing antibody responses, we developed a novel fluorescence-based VNT using our rOROVMZsG reporter virus^26^. This assay enabled rapid visualization of inhibition by monitoring ZsGreen expression in infected cells. Sera from pups born to rOROV-exposed dams exhibited strong, dose-dependent suppression of fluorescence. In contrast, sera from pups of naive dams showed little to no inhibition until 10 dpi (Fig. 6e, f). Neutralization titers derived from this assay demonstrate peak antibody activity at early time points, followed by a decline, consistent with transient, maternally derived protection (Fig. 6g; Exposed, red). To confirm these findings, we performed a complementary crystal violet-based VNT (Supplementary Fig. 4), which produced concordant results. Collectively, these data show that maternal OROV infection generates robust but short-lived neutralizing antibody-mediated protection in offspring.

## Discussion

Robust animal models are essential for elucidating the mechanisms of viral pathogenesis, transmission, and immunity in a physiologically relevant context^31–34^. Although OROV has historically been associated with self-limiting febrile illness, the 2022–2024 outbreak confirmed its capacity for maternal-to-fetal transmission. In this study, we combined *in vivo* and *in vitro* approaches to (a) establish a murine pregnancy model for OROV and (b) compare a contemporary isolate (OROV 240023 from the Cuban 2024 outbreak) with the ancestral strain (rOROV BeAn19991 isolated in 1960 from a Sloth in Brazil). Using timed gestational infections in immunocompetent C57BL/6J mice, we demonstrate that vertical transmission is a conserved feature of OROV biology. This finding underscores the importance of incorporating OROV into maternal–child health surveillance, diagnostic, and vaccine preparedness efforts.

Although vertical transmission potential was conserved, OROV 240023 showed clear signs of enhanced replicative fitness. In cell culture, OROV 240023 replicated faster than BeAn19991 in both IFN-competent and IFN-deficient cell lines, particularly during early infection, and in IFNAR^⁻/⁻^ mice, OROV 240023 caused earlier mortality. These kinetics likely contribute to the higher frequency of early gestation placental infection observed *in vivo*, even if end-stage tissue distribution appeared similar. This faster replication may provide a competitive advantage in natural transmission cycles, potentially affecting outbreak dynamics. To investigate underlying mechanisms, we performed minigenome assays with the contemporary Brazilian isolate. Here, AM0059/88 consistently outperformed BeAn19991 in reporter activity. While this enhancement is likely influenced by variation in the RdRp, we also identified mutations in the 3′ UTR of the M segment that may affect cis- or trans-acting regulating elements. As we have previously shown, mutations in OROV UTRs are crucial for minigenome activity^35^. Ongoing studies are focused on dissecting how these changes affect viral gene expression and virulence.

Molecular divergence was also observed in the glycoproteins. Scachetti *et al.*^30^ previously reported that human and murine sera raised against BeAn19991 poorly neutralized OROV AM0088, indicating antigenic mismatch. While our neutralization experiments were not epitope-resolved, we did observe that sera from BeAn19991-infected mice were less effective at neutralizing OROV 240023. Given that OROV 240023 and AM0059/88 share multiple conserved mutations within Gc, these strains likely exhibit similar antigenic profiles. This raises the possibility of immune evasion, whereby individuals previously exposed to antigenically distinct strains remain susceptible to reinfection, potentially contributing to the resurgence and geographic spread observed during the recent outbreak. Supporting this, prior work using vesicular stomatitis virus (VSV)-based OROV vaccine chimeras identified the N-terminal head domain of Gc as immunodominant but non-essential for protection^36^. Deletion of this variable domain redirected antibody response toward more conserved regions and improved protective efficacy, suggesting that natural infection may drive strain-specific immunity that limits cross-protection. Our sequence analysis confirms that contemporary strains harbor substitutions within this same region, which likely underlie reduced neutralization by ancestral-specific sera. Together, these findings support a model in which Gc variability facilitates immune escape, underscoring the importance of targeting conserved epitopes for vaccine design.

In addition to structural protein variation, changes in non-structural proteins NSm and NSs may also influence fitness and host interactions. We have previously shown that the NSm is dispensable for OROV in the mammalian host^26,27^, but its role in vector competence remains unresolved. Several variant residues are shared between 240023 and AM0059/88 and differ from BeAn19991, including substitutions that may potentially affect NSm function. Investigating the phenotypic consequences of these mutations will help determine whether NSm, as in other orthobunyaviruses, such as Bunyamwera virus, contributes to vector-specific replication or fitness^37^. Our findings, together with our recent publication^13^, show that OROV 240023 replicates more rapidly than the ancestral strain in both the mammalian host and the vector. Within the NSs, both 240023 and AM0059/88 encode an asparagine residue at position 71, while 240023 uniquely encodes a threonine residue at position 67. We have previously demonstrated that deletion of NSs results in robust type I IFN production, STAT1 phosphorylation, and upregulation of antiviral effectors such as MxA^27^. Because NSs overlaps the N gene in an alternative reading frame, such changes likely reflect evolutionary pressure to maintain N function while fine-tuning host immune antagonism. Whether these substitutions enhance NSs’ stability, IFN suppression, or host-specific interactions is currently under investigation.

While these molecular features shape viral replication and immune evasion, our study also uncovered important immunological consequences of OROV infection during pregnancy. We found that maternal OROV infection elicits neutralizing antibodies that are transferred to offspring, conferring partial protection upon homologous challenge. In our fluorescence-based VNT, sera from pups born to infected dams showed potent, dose-dependent inhibition of infection, which declined over time, consistent with transient, maternally derived protection, likely mediated by postnatal exposure through milk, given the timing of infection and delivery. These findings parallel what has been described for other vertically transmitted arboviruses, including Zika^38,39^ and dengue virus^40^, and highlight the potential for maternal immunization as a strategy to protect neonates in OROV-endemic regions.

While our C57BL/6J model provides a tractable and physiologically relevant system for studying OROV vertical transmission, it has limitations. Murine placental architecture differs from that of humans, and our results may under- or overestimate fetal susceptibility in human pregnancy. Infections were performed under controlled timing and inoculation routes, which may not fully capture the complexity of natural transmission. Fetal infection rates could also be influenced by gestational stage at exposure, maternal immunity, or viral dose; these variables warrant systematic study. Finally, the absence of overt fetal malformations in our model requires more detailed assessment.

In conclusion, we show that OROV can cross the placenta, infect the fetus, and induce maternal neutralizing antibodies that are transferred to offspring, providing partial protection from homologous challenge. These results provide direct experimental evidence of congenital OROV infection and demonstrate that vertical transmission is a conserved trait present in both ancestral and contemporary outbreak strains. Coupled with our mechanistic insights into viral replication, immune escape, and antigenic divergence, this work defines key biological properties that could influence OROV’s epidemic potential. Our murine and human trophoblast-based models, as well as a recently described human organoid model^41^, provide complementary platforms for dissecting how viral and host factors influence OROV vertical transmission and fetal outcomes. As OROV continues to expand its geographic and clinical footprint, understanding how viral evolution shapes transmission dynamics, tissue tropism, and immune evasion will be critical for guiding vaccine and therapeutic development. Given the high reassortment potential among orthobunyaviruses, including OROV itself^11,22,23,28,42^, even subtle sequence variation may have meaningful consequences for pathogenesis and antigenicity. Our findings emphasize the need for sustained genomic surveillance and functional virology to assess emerging OROV strains. Additionally, our work for the first time provides a suite of novel OROV reagents and models, spanning *in vitro* and *in vivo* systems, that will serve as valuable tools for studying OROV pathogenesis and evaluating vaccine candidates.

## Materials and Methods

### Ethics statement

All animal work was performed in compliance with Indiana University School of Medicine’s Institutional Animal Care and Use Committee (IACUC) under an approved IACUC protocol 22080 (PI: Tilston). All work was carried out in an Animal Biosafety Level 2 (ABSL-2) facility.

### Cells and viruses

Vero E6 cells (African green monkey kidney cells), A549 (human alveolar adenocarcinoma epithelial cells) and BSR-T7/5 cells^43^ (baby hamster kidney cells), which stably express T7 RNA polymerase were grown in Dulbecco’s modified Eagle medium (DMEM; Gibco) supplemented with 2 - 10% (V/V) fetal bovine serum (FBS; Gibco). BeWo cells (choriocarcinoma-derived trophoblasts) were grown in Kaighn’s Modification of Ham’s F-12 Medium (F-12K; ATCC) supplemented with 10% (V/V) FBS. JEG3 cells (choriocarcinoma-derived trophoblasts) were grown in Eagle’s minimal essential medium (EMEM; Sigma Aldrich) supplemented with 10% (V/V) FBS. HTR8 cells (HTR8/SVneo first-trimester transformed cells) were grown in RPMI-1640 supplemented with 10mM HEPES, 1mM sodium pyruvate, and 10% (V/V) FBS. All cells were grown at 37°C and 5% CO_2_.

We have previously described^26,27^ both rOROV^BeAn19991^ and rOROVMZsG. OROV^240023^ was obtained from the Arbovirus Reference Collection at the University of Texas Medical Branch (UTMB). For simplicity, throughout the manuscript and figures, rOROV^BeAn19991^ and OROV^240023^ are referred to by their strain designations, such as rOROV ancestral strain <u>BeAn19991</u> and OROV strain <u>2400023</u>, respectively. OROV 240023 was passaged 3 times in Vero E6 cells at UTMB and once upon arrival at our facility. To obtain pure stocks, we plaque-purified 240023. Here, infected Vero E6 cells were overlaid with 0.6% low-melt agarose (Fisher Scientific, BP16525) in 2 x minimum essential medium (MEM) with 2% FBS. 4 dpi cells were stained with 0.33% Neutral Red solution (Sigma Aldrich) to visualize plaques. Plaques were then picked using a pipette tip to infect Vero E6 cells (12-well plate, 2 x 10^5^ cells/ml). 48 hpi infectious virus supernatants were harvested. Virus stocks were grown and titrated in Vero E6 cells using a 50% Tissue Culture Infectious Dose Assay (TCID_50_). Virus titers for rOROV BeAn19991, rOROVMZsG, and OROV 240023 were 2.42×10^6^ TCID_50_/ml, 2.7×10^5^ TCID_50_/ml, and 1.1×10^7^ TCID_50_/ml, respectively.

### RT-PCR and Sanger Sequencing of OROV 240023

vRNA from OROV 240023 was extracted from 250 μl of viral supernatant using 750 μl of TRIzol LS Reagent (Ambion) and purified with the Direct-zol RNA MiniPrep kit (Zymo Research), following the manufacturer’s instructions. Complementary DNA (cDNA) synthesis was performed using M-MuLV reverse transcriptase (New England Biolabs). Segment-specific primers were used for PCR amplification of each genome segment: <u>S segment</u> (priOROVSF-FL: AGTAGTGTACTCCACAAT and priOROVSR-FL: AGTAGTGTGCTCCCAATT), <u>M segment</u> *fragment 1* (priOROV-MF-new-FL: AGTAGTGTACTACCAGCAACAA and priT7AM0059M4045R: GCAAAGCAGGTGATTTGTGTGC), <u>M segment</u> *fragment 2:* (priT7AM0059M2186F: GTAGATCCACCCGCTCAG and priOROV-MR-new-FL: AGTAGTGTGCTACCAACA), <u>L segment</u> *fragment 1* (priT7AM00590L91F: ATGTCGCAACTGTTACTC and priT7AM00590L4148R: GCTTAACAGGTATCTCTG), <u>L segment</u> *fragment 2* (priT7AM00590L2213F: CAGACTACGATATAACAC) and priT7AM00590L6328R: CTTATCATGTATGCTAGTG) and <u>L segment</u> *fragment 3* (priT7AM00590L3623F: CATCCATATGTATGGTGC and priT7AM00590L8546R: CTTAGAAGTCAAATTTGG). PCR amplification of the S segment was performed using GoTaq™ G2 Master Mix (Promega) in a thermal cycler (Applied Biosystems) with touchdown PCR (annealing temperatures: 45 °C, 48 °C, and 50 °C). M segment cDNA was amplified under standard PCR conditions with an annealing temperature of 56 °C. PCR products were resolved by agarose gel electrophoresis, and bands of the expected size were excised and purified using a gel extraction kit (IBI Scientific). Purified DNA was submitted for Sanger sequencing and analyzed using SeqMan Ultra, DNASTAR (V18.0.3.2).

### Viral growth kinetics

Vero E6 and A549 cells (2 x 10^5^cells/ml, 24-well plate) were infected with either rOROV BeAn19991 or OROV 230024 at an MOI 0.1 for 1 h at 37°C. Cell monolayers were then washed 3 x with D-PBS (Gibco) and then provided with a 2% FBS growth medium. At desired time points, supernatants were collected and harvested, and infectious viral titers determined by a TCID_50_ assay. HTR8, BeWo and JEG3 cells (1 x 10^5^ cells/ml, 24-well plate) were infected with either rOROV BeAn19991 or OROV 230024 at an MOI 0.002 for 1 h at 37°C. Cell monolayers were then washed 3 x with D-PBS (Gibco) and then provided with a 10% FBS growth medium. At desired time points, supernatants were collected and harvested, and a TCID_50_ assay determined infectious viral titers.

### TCID_50_ and plaque assays

TCID_50_ assays were performed in Vero E6 cells and seeded at a density of 10^4^ cells per well in 96-well plates and infected with a 10-fold serial dilution of virus. At 5 – 7 dpi, CPE was recorded, and viral titers were expressed as tissue culture infectious dose (TCID_50_ units). Plaque assays were performed in 6-well plates with Vero E6 cells at a density of 10^5^ cells/ml. Cell monolayers were infected with a 200 µl inoculum of virus diluted in Opti-MEM. Cells were then overlaid with 0.6% Avicel (FMC, Avicel RC-591) in 2 x minimum essential medium (MEM)/2% fetal bovine serum (FBS). Cells were fixed at 3 – 4 dpi with 4% (wt/vol) paraformaldehyde (PFA) in phosphate-buffered saline (PBS) for 30 min, and plaques were visualized using crystal violet.

### Minigenome assays

Sub-confluent BSR-T7/5 cell monolayers were seeded in 24-well plates at 1.4 x 10^5^ cells/ml and co-transfected with 500 ng of pT7-BeAn19991-M:hRen or pT7-AM0059/88-M:hRen reporter plasmid. For N protein titration experiments, cells were co-transfected with increasing amounts of pTM1-N (0-1000 ng) and constant amounts of pTM1-RdRp (250 ng). For RdRp titration experiments, cells received increasing amounts of pTM1-RdRp (0-1000 ng) with constant amounts of pTM1-N (250 ng). pTM1-FFLuc (Firefly luciferase; 5 ng) served as an internal normalization control, and pTM1-empty vector was added to maintain equal total DNA amounts across all transfections.

Transfection complexes were prepared by combining plasmid mixtures with 25 μl Opti-MEM (ThermoFisher Scientific) containing 1 μl P3000 enhancer reagent. Separately, lipid transfection mixtures were prepared in 25 μl Opti-MEM containing Lipofectamine 3000 reagent (0.6 μl per μg total DNA; ThermoFisher Scientific). After incubation and complex formation according to manufacturer’s protocol, transfection mixtures were added to cell monolayers. Cells were lysed 24 hours post-transfection (hpt), and *Renilla* and firefly luciferase activities were measured using the Dual-Luciferase Reporter Assay System (Promega) according to manufacturer’s instructions. *Renilla* luciferase values were normalized to firefly luciferase activity to account for transfection efficiency variations.

### Immunofluorescence

Confluent monolayers of Vero E6, HTR8, BeWo, or JEG3 cells were infected with either rOROV BeAn19991 or OROV 240023 at a set volume for virus isolation or an MOI of 0.1. After 24 h of incubation at 37°C, 5% CO_2_, the supernatant was discarded, and the cell monolayer was washed with phosphate-buffered saline (PBS) and fixed using 4% (w/v) paraformaldehyde in PBS. After 20 min of incubation at room temperature (RT), the fixative was removed, and the monolayer was washed twice with PBS. Cell monolayers were then permeabilized (0.5 % Triton X-100, 20 mM sucrose in PBS; 1 ml) for 30 min at RT. Cells were then incubated with 200 µl mouse anti-OROV diluted 1:500 in 1 % Bovine Serum Albumin (BSA)/0.1 % Tween-20 in PBS for 1 h at RT. Cells were washed several times before incubating with 200 µl Alexa Fluor 594 goat anti-mouse IgG (H&L) diluted 1: 1000 in 1 % BSA/0.1 % Tween-20 in PBS for 1 h at RT. After washing the cells three times with PBS, 4′,6-diamidino-2phenylindole (DAPI) nuclear stain (Fisher EN2248) was added and incubated in the dark for 10 minutes at RT. BeWo cells were incubated with 200 µl Alexa Fluor 488 phalloidin (Invitrogen A12379) diluted 1: 500 in PBS for 15 min at RT to stain cellular actin. Images were obtained using a fluorescent microscope (EVOS M5000 imaging system).

### Animal study design

a. <u>Pathogenesis study in C57BL/6J mice</u>: Six-week-old female C57BL/6J mice (Jackson Laboratory, IUSM colony) were housed under specific pathogen-free conditions in HEPA-filtered cages within an ABSL-2 facility, with ad libitum access to food and water. Mice were subcutaneously (SC) inoculated with 10^5^ TCID_50_ of rOROVMZsG diluted in Opti-MEM to a final volume of 100 μl. Control animals received UV-inactivated rOROVMZsG (Benchmark UV Clave chamber; 6 × 4-W UV bulbs; 150 mJ/cm^2^) at the same dose and volume. Mice were monitored daily for clinical signs, and body weights were recorded. On 5, 7, and 14 dpi, animals were anesthetized with isoflurane and terminally exsanguinated for serum collection, followed by euthanasia via cervical dislocation. Tissues, including liver, spleen, heart, lungs, and brain, were collected for viral isolation and RNA extraction. For viral load analysis, organs were placed in pre-weighed tubes containing 500 μl PBS supplemented with 2 x Antibiotic-Antimycotic (Gibco), weighed, and homogenized using a Bead Mill 24 Homogenizer (Fisherbrand). A 100 μl aliquot of homogenate was mixed with 400 μl TRIzol LS reagent (Ambion) and processed using the Direct-zol RNA Purification Kit (Zymo Research). (b) <u>Timed pregnancy studies:</u> Six-week-old female C57BL/6J mice (Jackson Laboratory, IUSM colony) were time-mated and housed under the conditions described above. The presence of a vaginal plug confirmed pregnancy, and at the indicated embryonic days, pregnant dams were subcutaneously inoculated with 2.42×10^5^ TCID_50_/ml of rOROV BeAn19991. Mice were monitored daily and weighed until euthanasia. Dams were anesthetized with isoflurane and terminally exsanguinated, followed by cervical dislocation. Maternal tissues (liver, spleen), placentas, and pups were collected for viral isolation and RT-qPCR analysis. Pups were euthanized by decapitation under ice water anesthesia or fixed in 10% buffered formalin (Fisher) for histological analysis. Maternal organs were processed as above using 500 μl PBS with 4x Antibiotic-Antimycotic. For pup viral loads, whole pups were collected in pre-weighed tubes containing 3 ml PBS with 4x Antibiotic-Antimycotic, weighed, homogenized, and 250 μl of homogenate was mixed with 750 μl TRIzol LS for RNA extraction. (c) <u>Pathogenesis</u> study in IFNAR^-/-^ mice: Six-week-old male and female IFNAR^⁻/⁻^ mice (Jackson Laboratory, IUSM colony) were housed in the ABSL-2 facility as described above. Mice were SC infected with 10^5^ TCID_50_/ml of either rOROV BeAn19991 or OROV 240023 diluted in Opti-MEM (100 μl total volume). Animals were monitored daily for clinical signs and weighed. Euthanasia was performed according to a predefined clinical scoring system as previously described^26^. Surviving animals were rechallenged with the same virus at 14 dpi. At euthanasia, mice were anesthetized and terminally exsanguinated. Tissues (liver, spleen, heart, lungs, brain) were collected for RT-qPCR and fluorescence analysis using an EVOS M5000 imaging system (ThermoFisher). For histology, tissues were fixed in 4% PFA, paraffin-embedded, and sectioned (5 μm) for hematoxylin and eosin (H&E) staining by the Indiana CTSI Histology Core.

### RT-qPCR assay

rOROV BeAn1991 vRNA was quantified using primers targeting the S genome segment as previously described^26^. OROV 240023 vRNA was quantified targeting the S genome segment using the following primers and probe: OROV-TVP-S qPCR primers (Fwd: TACATCGCGTCACCATCATTC, Rev: CCCAGATGCGATCACCTATTAAG, OROV-TVP-SqPCRprobe FAM: TTGGCAGAGGTGAAGGGTTGTACT). The working concentration for the primers was 10 μM for the probe 5 μM. To generate an OROV 240023 S RNA standard curve, an IDT gene fragment containing the sequenced OROV 240023 S segment was synthesized and used as a template for *in vitro* RNA transcription (MEGAscript T7 transcription kit; Invitrogen). The resulting RNA was diluted to known copies per ml in RNase-free water and serially diluted for each assay. RT-qPCR was performed using the Luna Universal Probe One-Step RT-qPCR Kit (New England Biolabs). The assay was performed on the QuantStudio^™^ 5 (ThermoFisher Scientific) using the following conditions: 55°C for 10 min, 95°C for 1 min, and then 40 cycles of 95°C for 10 sec and 60°C for 1 min. For the animal studies, RNA levels were reported as log vRNA copies per gram of tissue, with the limit of detection calculated based on the highest cycle threshold (CT) value detected in the standard curve for each assay.

### Hybridization Chain Reaction (HCR) for OROV vRNA detection in tissue sections

Probe sets targeting rOROV BeAn19991 S segment were designed using the DIY probe generator pipeline^44^. All probes were screened for specificity against host cell transcripts to minimize background. The HCR assay was performed following protocols published by Molecular Instruments, Inc. To visualize OROV vRNA, tissue sections were deparaffinized in Histoclear (3x 5 min), followed by two washes in 100% ethanol (3 min each) to remove residual Histoclear. Sections were rehydrated using a graded ethanol series (95%, 85%, 70%, and 50% ethanol, each for 3 min) and rinsed in Milli-Q water for 3 min. For antigen retrieval, slides were immersed in 250 ml of 1x antigen unmasking solution (prepared from a 100× stock in Milli-Q water) within a plastic container, ensuring complete submersion. The container was placed in a pressure cooker and heated at maximum pressure for 20 min. After retrieval, slides were cooled to RT and incubated in 1x PBST (2x 2 min). Tissue sections were then treated with 10 µg/ml proteinase K in PBST for 10 min at 37°C in a humidified chamber, followed by two washes in fresh 1x PBST. Probe hybridization buffer (50–100 µl per section) was applied, and slides were incubated at 37°C for 10 min in a humidified chamber. Hybridization was performed by adding a probe solution (0.4 pmol of each probe set; Supplemental Table; in 100 µL of hybridization buffer), followed by overnight incubation (>12 h) at 37°C. The next day, slides were washed sequentially in probe wash buffer at 37°C for 15 min, followed by graded washes in 75%, 50%, and 25% probe wash buffer diluted in 5x SSCT, and finally 100% 5x SSCT (15 min per step). Slides were then immersed in 5x SSCT at RT for 5 min before air-drying. For amplification, sections were incubated with an amplification buffer at RT for 30 min. Hairpin reagents (B3-647; Molecular Instruments, Inc.) were prepared by snap cooling 6 pmol of hairpins h1 and h2 separately (2 µl of 3 µM stock, heated at 95°C for 90 sec, then cooled to RT for 30 min in a dark tube). The snap-cooled hairpins were then mixed in 100 µl of amplification buffer. The pre-amplification solution was removed, and 50–100 µl of the hairpin solution was applied to each section, followed by overnight incubation (>12 h) in a dark, humidified chamber at RT. On the final day, slides were washed four times for 5 min each in 5x SSCT at RT. Nuclei were stained with 100 µl of 1 µg/ml Hoechst for 5 min, followed by a final wash in 5x SSCT. Sections were immediately mounted and visualized on the EVOS M5000 imaging system (ThermoFisher).

### Virus neutralization assays

a. <u>High-throughput assay using reporter virus:</u> Mouse sera was heat-inactivated at 56 °C for 30 min in a thermal cycler (Applied Biosystems), then 2-fold serially diluted in FluoroBrite™ DMEM (Gibco) in 96-well dilution plates (CellTreat; 25 μl/well). Diluted rOROVMZsG (MOI 0.1) was added to each well and incubated at 37 °C for 1 h. Virus–serum mixtures were transferred to Vero E6 cells (10^4^ cells/well) seeded in black-walled 96-well plates (Corning). Sera from a Lone Star virus (LSV)-infected mouse served as a negative control. Plates were incubated for 48 h and read using a BioTek Synergy H1 plate reader. Whole-plate fluorescent images were acquired using an Odyssey M Imager (LI-COR). (b) <u>Neutralization of ancestral and contemporary strains:</u> Sera from rOROV BeAn19991-infected mice were heat-inactivated and serially diluted 2-fold in Opti-MEM (25 μl/well). Diluted rOROV BeAn19991 or OROV 240023 (MOI 0.1) was added and incubated for 1 h at 37 °C before transfer to Vero E6 cells (10^4^ cells/well in 96-well plates). At 5 dpi, cells were fixed in 4% PFA (15 min) and stained with crystal violet. Neutralization titers were defined as the highest serum dilution exhibiting <50% CPE.

### Data and statistical analysis

RT-qPCR data were processed in Microsoft Excel and visualized using GraphPad Prism (V10). vRNA levels between OROV BeAn19991 and OROV 240023 were analyzed using Mann-Whitney analysis. Χ² analysis was used to compare rates of placental infection between strains. (Fig. 1m). Growth comparison of rOROV BeAn19991 and OROV 240023 was analyzed by a two-way ANOVA with Sidak’s multiple comparisons test (Fig. 3a) and a multiple unpaired t-test (Fig. 2a-c) with individual variance, with false discovery rate (FDR) correction applied using the two-stage step-up method of Benjamini, Krieger, and Yekutieli in GraphPad Prism (V10). Sliding window analysis and visualization were performed in R (V4.4.0) and RStudio (V2024.04.1+682) (Fig. 3d). Sequencing alignments and visualization were performed using DNAstar (V18.0.3.2) (Fig. 3e and 5e).

## Data availability

All relevant data are included within the manuscript and its supporting files. Additional information can be obtained from the corresponding author (N.L.T.).

## Acknowledgments

We thank Dr. Andrew M. Lunel (Department of Pediatrics, Indiana University School of Medicine, IUSM) for advice on tissue processing, and Lylah Hutson and Dr. George Sandusky (Department of Pathology, IUSM) for histology consultation. We are grateful to Drs. William M. de Souza (University of Kentucky, U.S), José Luiz Proenca-Modena (University of Campinas, Brazil), and Pritesh Lalwani (Instituto Leônidas e Maria Deane, Fiocruz Amazônia, Brazil) for generously providing genome sequences for Brazilian OROV isolates before they became publicly available. We thank the Laboratory Animal Resource Center (LARC) staff at ISUM for assisting with mouse husbandry and care throughout this study. This work was supported in the United States by a Showalter Trust award, NIH grant R03AI190651, and IUSM start-up funds to N.L.T. In the United Kingdom, A.T.C., M.McF., and B.B. were supported by a Wellcome Trust/Royal Society Sir Henry Dale Fellowship (210462/Z/18/Z) awarded to B.B., and by the MRC Core Award (MC_UU_00034/4).

## Author contributions

N.L.T. and K.G. conceptualized the study. N.L.T., K.G., J.V., and B.B. designed the experiments. K.G., J.B., A.T.C., M. McF., D.C.A.O., S.P., and N.L.T. performed the experiments. D.A. and L.R. contributed knowledge and reagents towards the human placental cell lines. N.L.T., K.G., J.B., J.V., and B.B. wrote the paper. All authors read and approved the manuscript for publication.

## Supplemental Data

**Supplemental Figure 1.**
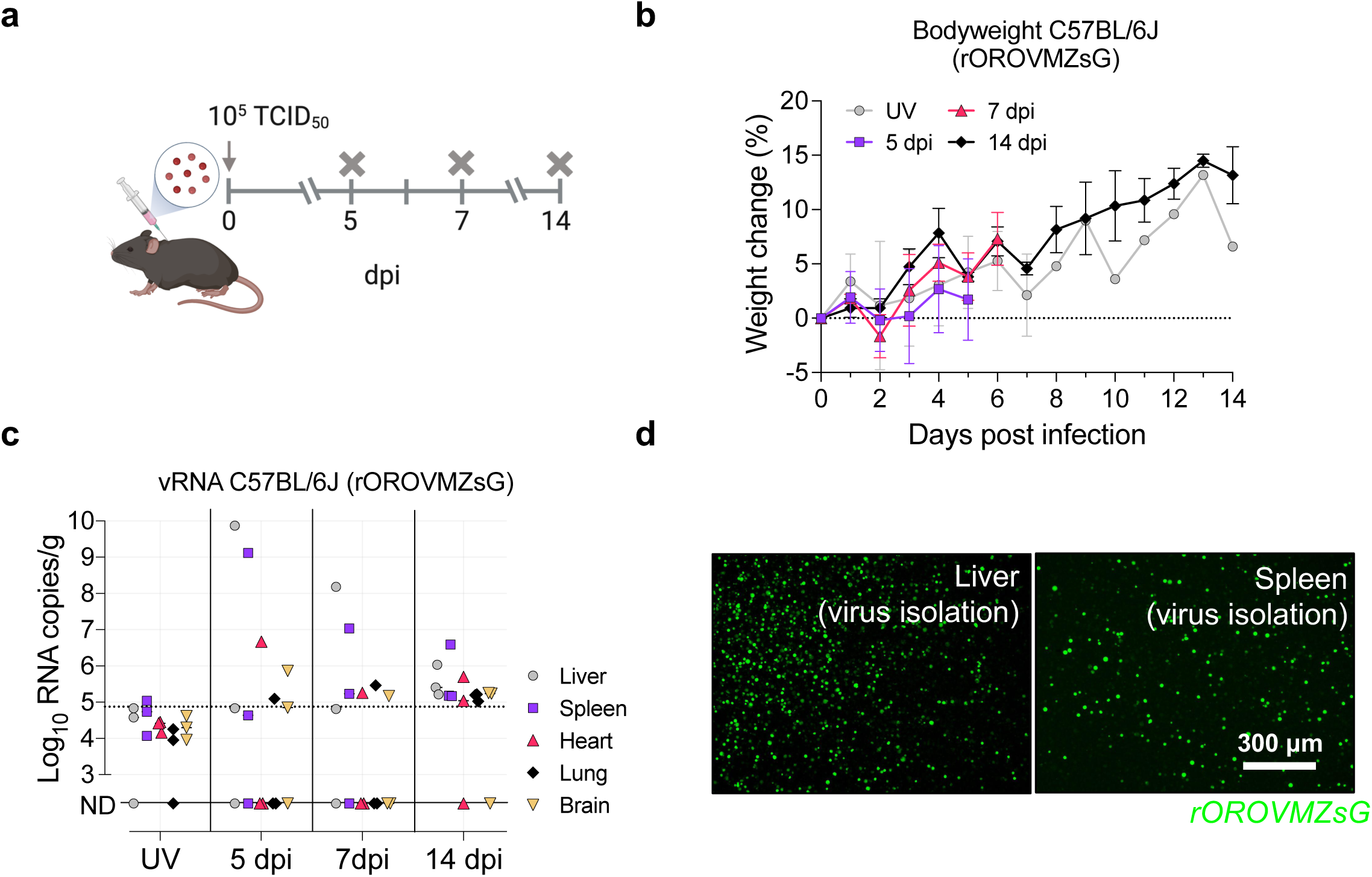
rOROVMZsG in immunocompetent C57BL/6J mice. (a) Schematic of experimental design. WT C57BL/6J mice were infected SC with either rOROVMZsG (3 groups, each n=3) or UV-inactivated rOROVMZsG (n=3), and euthanized at 5, 7, or 14 dpi. (b) Percent weight change of mice from baseline compared to UV-inactivated virus controls. (c) vRNA loads per gram of tissue in the liver, spleen, heart, lung, and brain of rOROVMZsG or UV-inactivated infected mice as measured by RT-qPCR. The dashed line represents the limit of detection, ND = not detected. (d) Representative fluorescence images from virus isolations in Vero E6 cells showing infectious rOROVMZsG in homogenized liver and spleen samples from infected mice (EVOS M5000 imaging system, ThermoFisher).

**Supplemental Figure 2.**
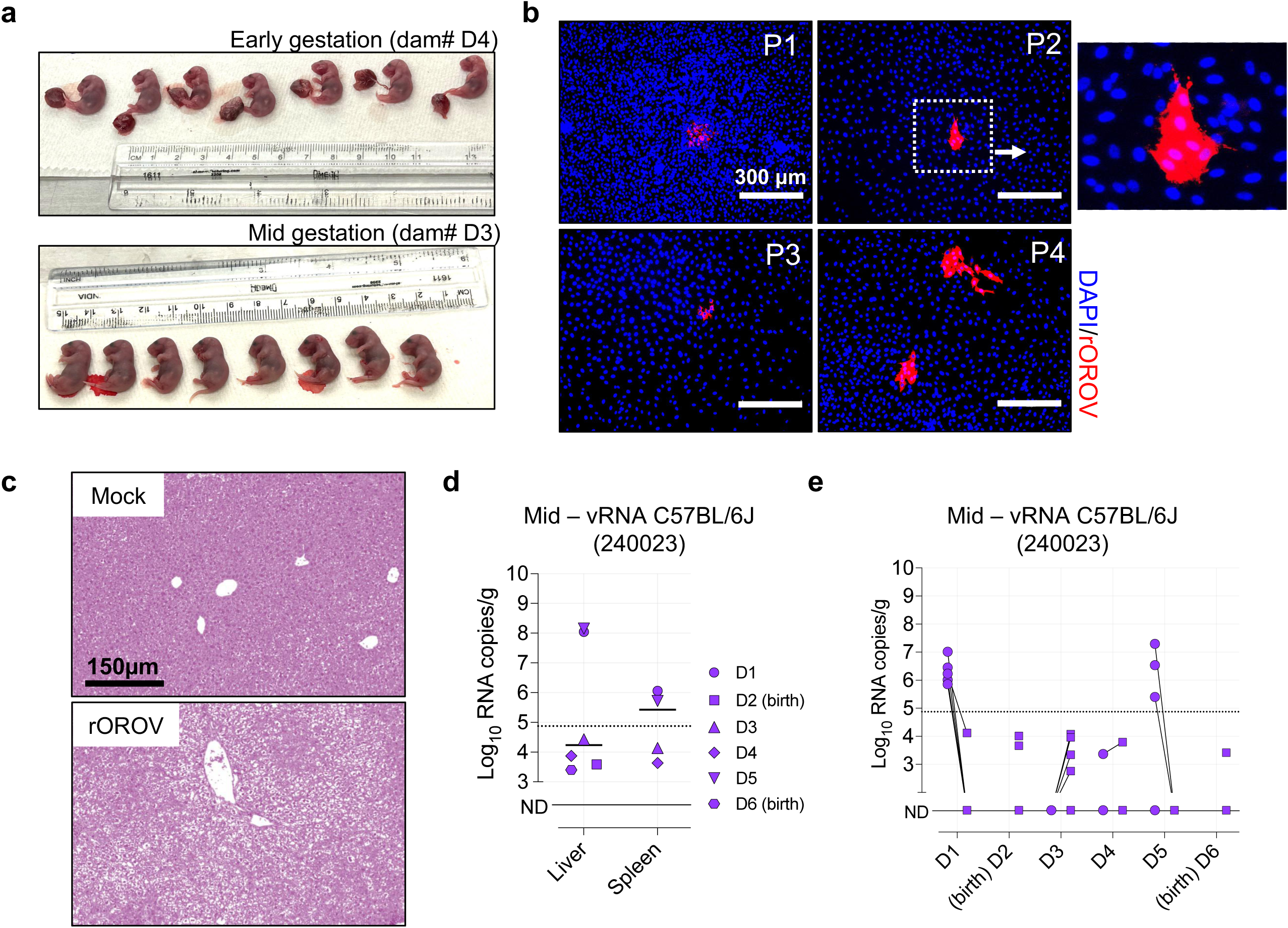
Infection with rOROV BeAn19991 causes liver pathology and placental infection. (a) Representative images of fetuses harvested from early or mid-gestation infected dams, showing normal gross morphology. (b) Detection of replication-competent virus in Vero E6 cells infected with placental homogenates. Immunofluorescence staining shows rOROV (red) at 24 hpi. Nuclei stained with DAPI (blue). Four representative placental isolates (#P1–P4) are shown. (c) Representative H&E-stained liver sections from mock-and rOROV-infected pregnant C57BL/6J dams. (d) vRNA levels in the liver and spleen of pregnant mice infected at mid-gestation with OROV 240023 (e) vRNA levels in matched placentas and fetuses following mid-gestation infection. The dashed line represents the limit of detection. ND = not detected.

**Supplemental Figure 3.**
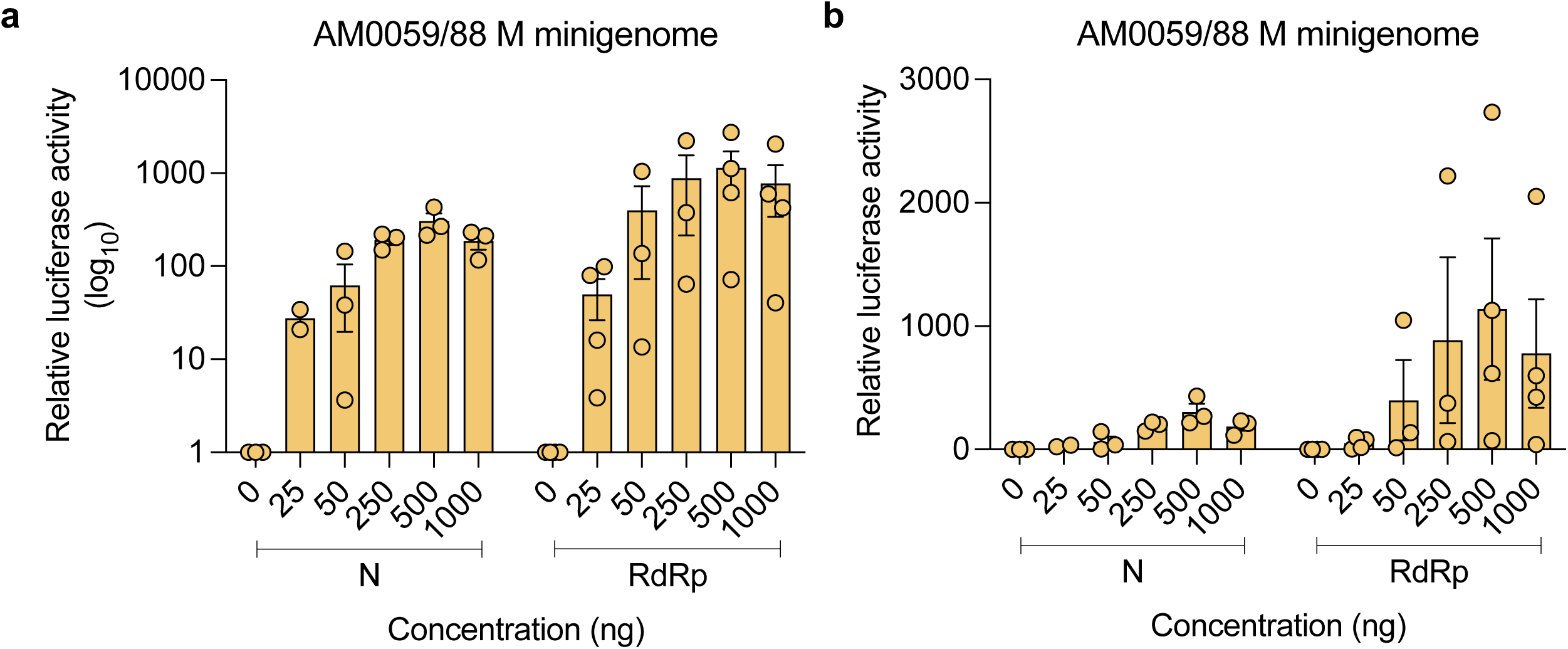
Optimization of AM0059/88 M segment minigenome assay. BSRT7/5 cells were co-transfected with either increasing amounts of pTM1-N and constant amounts of pTM1-RdRp (250 ng) or increasing amounts of pTM1-RdRp and constant amounts of pTM1-N (250 ng), along with an AM0059/88 M-segment minigenome encoding humanized Renilla luciferase (hRenilla). A firefly luciferase-expressing plasmid, pTM1-FFLuc was also included in the transfection mixture to normalize for differences in transfection efficiency. Data are shown as relative luciferase activity (a) log10 scale and (b) linear scale. Bars represent the mean of biological replicates, with individual data points shown.

**Supplemental Figure 4.**
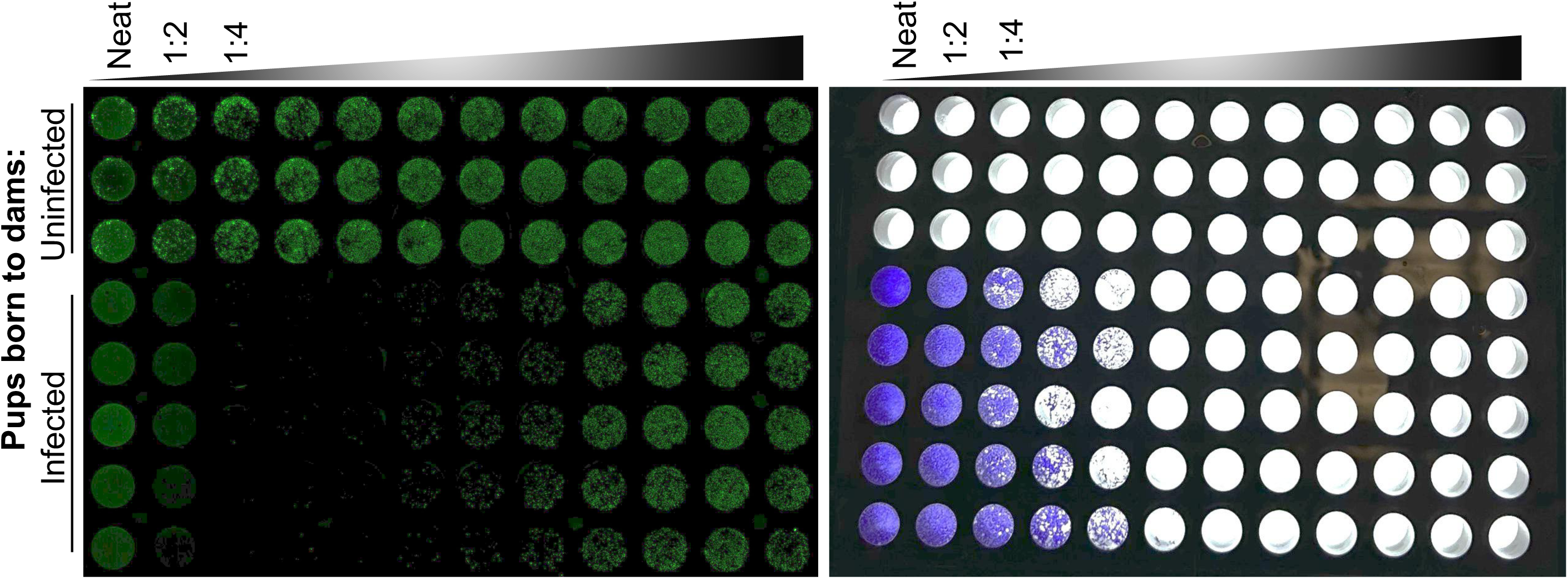
High-throughput VNT assay set-up. High-throughput VNT performed in a 96-well format using the same sera as in Fig. 6e. LI-COR fluorescence scans (left) and crystal violet-stained plates (right) show a dose-dependent viral inhibition in samples from pups born to infected dams. Representative data shown from 3 dpi.

---

**Supplemental Table:**
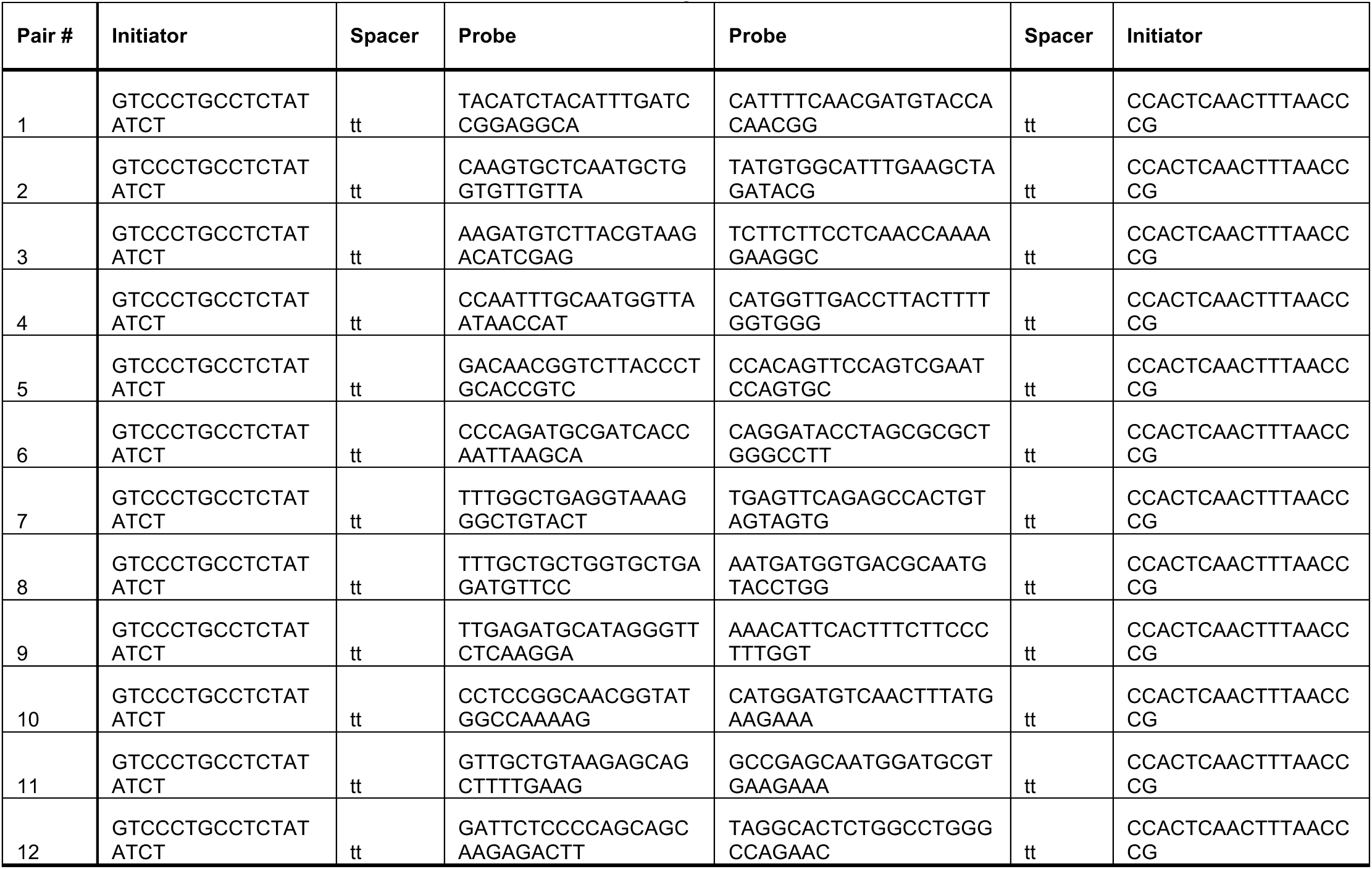
HCR probes used in this study.

## Notes

### Competing Interest Statement

The authors have declared no competing interest.

## References

1. PAHO/WHO. Epidemiological Update Oropouche in the Region of the Americas, 11 February 2025. (Pan American Health Organization, World Health Organization, 2025).

2. Salvato, R.S. Re-emergence of Oropouche virus as a novel global threat. Current Research in Microbial Sciences 8, 100406 (2025).

3. Naveca, F.G., et al. Human outbreaks of a novel reassortant Oropouche virus in the Brazilian Amazon region. Nature Medicine 30, 3509–3521 (2024).

4. PAHO/WHO. Epidemiological Alert Oropouche in the Region of the Americas, 6 September 2024. (Pan American Health Organization, World Health Organization, 2024).

5. PAHO/WHO. Epidemiological Alert Oropouche in the Region of the Americas: vertical transmission event under investigation in Brazil, 17 July 2024. (Pan American Health Organization, World Health Organization, 2024).

6. PAHO/WHO. Epidemiological Update Oropouche in the Region of the Americas, 12 April 2024. (Pan American Health Organization, World Health Organization, 2024).

7. PAHO/WHO. Epidemiological Alert Oropouche in the Region of the Americas, 9 May 2024. (Pan American Health Organization, World Health Organization, 2024).

8. PAHO/WHO. Epidemiological Update Oropouche in the Region of the Americas, 6 March 2024. (Pan American Health Organization, World Health Organization, 2024).

9. PAHO/WHO. Epidemiological Alert Oropouche in the Region of the Americas, 2 February 2024. (Pan American Health Organization, World Health Organization 2024).

10. PAHO/WHO. Epidemiological Alert Oropouche in the Region of the Americas, 1 Aug 2024. (Pan American Health Organization, World Health Organization, 2024).

11. Tilston-Lunel, N.L. Oropouche Virus: An Emerging Orthobunyavirus. Journal of General Virology 105(2024).

12. Bello-Rodríguez, B.M., Vega-Jiménez, J., Cañete, R. & Rodriguez-Morales, A.J. Emergence of Oropouche virus infection in Matanzas, Cuba, 2024. Journal of Infection 90(2025).

13. Scroggs, S.L.P., et al. Enhanced infection and transmission of the 2022–2024 Oropouche virus strain in the North American biting midge Culicoides sonorensis. Scientific Reports 15, 27368 (2025).

14. Tilston, N., Skelly, C. & Weinstein, P. Pan-European Chikungunya surveillance: designing risk stratified surveillance zones. Int J Health Geogr 8, 61 (2009).

15. Elliott, R.M.> Bunyaviruses and climate change. Clinical Microbiology and Infection 15, 510–517 (2009).

16. das Neves Martins,> F.E., et al. Newborns with microcephaly in Brazil and potential vertical transmission of Oropouche virus: a case series. The Lancet Infectious Diseases 25, 155–165 (2025).

17. Ribeiro, B.F.R., et al. Congenital Oropouche in Humans: Clinical Characterization of a Possible New Teratogenic Syndrome. Viruses 17(2025).

18. Cola, J.P., et al. Maternal and Fetal Implications of Oropouche Fever, Espírito Santo State, Brazil, 2024. Emerg Infect Dis 31, 645–651 (2025).

19. Bandeira, A.C., et al. Fatal Oropouche Virus Infections in Nonendemic Region, Brazil, 2024. Emerg Infect Dis 30, 2370–2374 (2024).

20. de Souza, W.M., et al. ICTV Virus Taxonomy Profile: Peribunyaviridae 2024. J Gen Virol 105(2024).

21. Elliott, R.M. Orthobunyaviruses: recent genetic and structural insights. Nature Publishing Group 12(2014).

22. Briese, T., Calisher, C.H. & Higgs, S. Viruses of the family Bunyaviridae: Are all available isolates reassortants? Virology 446, 207–216 (2013).

23. Tilston-Lunel, N.L., et al. Genetic analysis of members of the species Oropouche virus and identification of a novel M segment sequence. Journal of General Virology 96, 1636–1650 (2015).

24. Gerrard, S.R., Li, L., Barrett, A.D. & Nichol, S.T. Ngari Virus Is a Bunyamwera Virus Reassortant That Can Be Associated with Large Outbreaks of Hemorrhagic Fever in Africa. Journal of Virology 78, 8922–8926 (2004).

25. Briese, T., Bird, B., Kapoor, V., Nichol, S.T. & Lipkin, W.I. Batai and Ngari Viruses: M Segment Reassortment and Association with Severe Febrile Disease Outbreaks in East Africa. Journal of Virology 80, 5627–5630 (2006).

26. Gunter, K., et al. A reporter Oropouche virus expressing ZsGreen from the M segment enables pathogenesis studies in mice. J Virol, e0089324 (2024).

27. Tilston-Lunel, N.L., Acrani, G.O., Randall, R.E. & Elliott, R.M. Generation of Recombinant Oropouche Viruses Lacking the Nonstructural Protein NSm or NSs. Journal of Virology 90, 2616–2627 (2016).

28. Bowen, J.M., et al. Probing orthobunyavirus reassortment using Bunyamwera and Batai viruses as models. PLoS Negl Trop Dis 19, e0013120 (2025).

29. Recaioglu, H. & Kolk, S.M. Developing brain under renewed attack: viral infection during pregnancy. Front Neurosci 17, 1119943 (2023).

30. Scachetti, G.C., et al. Reemergence of Oropouche virus between 2023 and 2024 in Brazil. medRxiv, 2024.2007.2027.24310296 (2024).

31. Shi, Y., et al. Vertical Transmission of the Zika Virus Causes Neurological Disorders in Mouse Offspring. Scientific Reports 8, 3541 (2018).

32. Roark, H.K., Jenks, J.A., Permar, S.R. & Schleiss, M.R. Animal Models of Congenital Cytomegalovirus Transmission: Implications for Vaccine Development. The Journal of Infectious Diseases 221, S60–S73 (2020).

33. Yadav, K.K. & Kenney, S.P. Animal Models for Studying Congenital Transmission of Hepatitis E Virus. Microorganisms 11(2023).

34. Morrison, T.E. & Diamond, M.S. Animal Models of Zika Virus Infection, Pathogenesis, and Immunity. Journal of Virology 91, 10.1128/jvi.00009-00017 (2017).

35. Acrani, G.O., et al. Establishment of a minigenome system for Oropouche virus reveals the S genome segment to be significantly longer than reported previously. Journal of General Virology 96, 513–523 (2015).

36. Stubbs, S.H., et al. Vesicular Stomatitis Virus Chimeras Expressing the Oropouche Virus Glycoproteins Elicit Protective Immune Responses in Mice. mBio 12, e0046321 (2021).

37. Terhzaz, S., et al. NSm is a critical determinant for bunyavirus transmission between vertebrate and mosquito hosts. Nat Commun 16, 1214 (2025).

38. Shan, C., et al. Maternal vaccination and protective immunity against Zika virus vertical transmission. Nature Communications 10, 5677 (2019).

39. Kim, I.J., et al. Zika purified inactivated virus (ZPIV) vaccine reduced vertical transmission in pregnant immunocompetent mice. NPJ Vaccines 9, 32 (2024).

40. Lee, P.X., Ong, L.C., Libau, E.A. & Alonso, S. Relative Contribution of Dengue IgG Antibodies Acquired during Gestation or Breastfeeding in Mediating Dengue Disease Enhancement and Protection in Type I Interferon Receptor-Deficient Mice. PLoS Negl Trop Dis 10, e0004805 (2016).

41. Megli, C.J., et al. Oropouche virus infects human trophoblasts and placenta explants. Nature Communications 16, 6040 (2025).

42. Tilston-Lunel, N.L., Shi, X., Elliott, R.M. & Acrani, G.O. The potential for reassortment between oropouche and schmallenberg orthobunyaviruses. Viruses 9, 1–12 (2017).

43. Buchholz, U.J., Finke, S. & Conzelmann, K.-K. Generation of Bovine Respiratory Syncytial Virus (BRSV) from cDNA: BRSV NS2 Is Not Essential for Virus Replication in Tissue Culture, and the Human RSV Leader Region Acts as a Functional BRSV Genome Promoter. Journal of Virology 73, 251–259 (1999).

44. Kuehn, E., et al. Segment number threshold determines juvenile onset of germline cluster expansion in Platynereis dumerilii. J Exp Zool B Mol Dev Evol 338, 225–240 (2022).

